# SRY-Box Transcription Factor 9 regulates 3’-Phosphoadenosine 5’-Phosphosulfate Synthase 2 mRNA expression through derepression of the transcriptional repressor, CCAAT/enhancer-binding protein beta

**DOI:** 10.1101/2024.09.11.612485

**Authors:** Cunren Liu, Rosa Serra

## Abstract

Previously, we showed that *Papss2* expression is regulated through a Sox9-dependent pathway. Here we explore molecular mechanisms whereby Sox9 regulates mouse *Papss2*. A 509bp Sox9-responsive DNA element (Region C) was identified upstream of the *Papss2* second start site using co-transfection and luciferase reporter assays. A Sox9 responsive element was narrowed down to 32bps within Region C (Sox9RE). Putative SoxE and C/EBPβ binding sites were identified within S9RE. C/EBPβ was identified as a repressor for Sox9-mediated activity. In cells transfected with expression vectors for C/EBPβ and Sox9, increasing amounts of C/EBPβ resulted in attenuation of Sox9-mediated activation of Region C while increasing amounts of Sox9 activated transcription in the presence of C/EBPβ. Using electromobility shift assays, three protein complexes were identified on S9RE after incubation with nuclear extracts from ATDC5 cells. Super shift assays indicated that under basal conditions C/EBPβ was present in the DNA-protein complexes observed. Unlabeled S9RE with point mutations in the predicted SoxE binding site competed with protein complex formation on the S9RE while excess oligo corresponding to the predicted SoxE binding site did not, suggesting that proteins do not bind to SoxE motiff under basal conditions. Under conditions of high Sox9 expression, the formation of protein-DNA complexes on S9RE was inhibited. We then showed by western blot that increasing Sox9 protein resulted in reduced C/EBPβ protein levels. Co-immunoprecipitation indicated interaction of Sox9 and C/EBPβ proteins. We propose that Sox9 acts to derepress C/EBPβ-inhibited transcription of Papss2 by first interacting with C/EBPβ to prevent it from binding DNA, then reducing C/EBPβ expression.

## Introduction

Papss2 is a bifunctional enzyme whose main function is to generate PAPS, the sulfate donor for sulfotransferase reactions (1, 2). Sulfation is especially important in articular cartilage where the large amount of sulfation on the glycosamino glycans linked to the core proteins of proteoglycans provide the necessary biomechanical properties for cartilage to function. Mutations in human PAPSS2 are associated with Pakistani type spondyloepimetaphyseal dysplasia (OMIM 612847) and a spontaneous mutation in mice, brachymorphic (bm), results from an autosomal recessive mutation in the Papss2 gene (3, 4). Papss2 is highly expressed in cartilage (3, 5) and the importance of this enzyme in cartilage biology is clear from the cartilage phenotypes associated with mutations in this gene (3).

Very little is known about how the levels of this enzyme are regulated even though matrix sulfation is essential for the maintenance of articular cartilage. The transcription factor Sex Determining region Y-box 9 (Sox9), along with Sox5 and Sox6, play an important role in the commitment of mesenchymal cells toward the chondrocyte lineage (6). Sox9 is member of the SoxE subgroup within the Sox family of HMG-box type DNA binding proteins, along with Sox8 and Sox10 (7) and Sox8 and Sox9 have some overlapping activities (8). Sox9 is expressed in mature articular chondrocytes and high expression levels of Sox9 are associated with maintenance of articular cartilage (9). It was previously shown that Papss2 and Sox9 are co-expressed during skeletal development supporting a potential role for Sox9 in mediating Papss2 expression (5). We previously showed that Sox9 was sufficient to up-regulate Papss2 mRNA whereas Sox9 did not appear to regulate another cartilage enriched gene, Prg4 (10). In addition, it was shown that Sox9 is required for up-regulation of Papss2 by TGFβ (10–12).

CCAAT/Enhancer-binding Proteins (C/EBP) are a family of transcription factors that containing a DNA binding domain and a leucine zipper dimerization domain. C/EBPβ has been shown to repress cartilage enriched genes including Cd-rap and Col2a1 (13–16). Expression of Cd-rap mRNA in cartilage involves derepression of C/EBPβ through its binding and sequestration by CREB-binding protein/p300 (CBP/p300) (13, 15). Sox9 bound to paired binding sites downstream of the C/EBPβ site, which were activated by CBP/P300 (13, 15). In contrast, C/EBPβ was shown to act as a transcriptional activator for genes enriched in hypertrophic chondrocytes including MMP13, Col10a1, and Runx2 (16, 17). It was suggested that C/EBPβ plays an important role in regulating the pre-hypertrophic to hypertrophic transition in chondrocytes.

In this study, we sought to determine the molecular mechanisms that regulate mouse Papss2 mRNA expression using the ATDC5 chondrogenic cell line. We identified a 32-bp Sox9 responsive element (S9RE) that contained predicted binding motifs for SoxE and C/EBPβ transcription factors. Increased C/EBPβ inhibited Sox9-mediated S9RE activity in a dose-dependent manner in co-transfection/reporter assays while excess Sox9 resulted in activation in the presence of C/EBPβ. EMSA and super shift assays indicated that C/EBPβ bound to S9RE under basal conditions when Papss2 expression levels were low. EMSA competition assays suggested that the SoxE motiff was not necessary for the formation of these DNA-protein complexes. When cells were infected with adenovirus vectors that express Sox9 and induce Papss2 expression, DNA-protein complex formation on S9RE was inhibited and C/EBPβ expression was down-regulated. In addition, co-immunoprecipitation assays demonstrated interaction of Sox9 and C/EBPβ proteins. We propose that Sox9 regulates Papss2 mRNA expression through derepression of C/EBPβ-mediated transcriptional repression on the S9RE by both down-regulating C/EBPβ protein levels and interacting with C/EBPβ, preventing it from binding to the DNA.

## Results

### Identification of Sox9 responsive elements (S9RE) in the Papss2 gene

To begin to identify molecular mechanisms whereby Sox9 regulates Papss2, we cloned several large evolutionarily conserved regions in and around the *Papss2* gene upstream of a 95bp minimum Col2a1 promoter luciferase reporter vector (18, 19) and tested their responsiveness to Sox9 using co-transfection followed by luciferase assays (Figure 1). We identified two Sox9 responsive regions, Region C and D (Figure 1B). Region C was located in the first intron between the first and second Papss2 transcriptional start sites corresponding to mouse locus GRCmm39 19:32,583,676-32,584,157 (Figure 1A). Regions D was located upstream of the first start site, GRCmm39 19:32,532,652-32,533,269. Region B was also in the first intron (GRCmm39 19:32,582,187-32,582,427) and was not responsive to Sox9. Region C and D contained previously identified Sox9 ChiP seq peaks from cartilage tissues (20, 21) and Figure 2A). This study will focus on Region C. To narrow down the DNA within Region C containing the Sox9 responsive element, fragments of the 509bp region were generated and multimers of the fragments were placed in luciferase vectors. Co-transfection/ luciferase assays were performed until a 32bp DNA fragment (GRCmm39 19:32,583,962-32,583,993) containing a functional Sox9 response element was identified (S9RE, Figure 2A, B). A predicted paired SoxE binding site (prediction score 270) was identified in the 32bp S9RE using the Jaspar 2024 database in the UCSC genome browser (Figure 2D; (22). A C/EBPβ binding site (prediction score 322) was also identified (Figure 2D). Mutations in the predicted SoxE binding site within the 32bp S9RE (Figure 2B) and within Region C (Figure 2C, highlighted red in Figure 2D) attenuated luciferase activity in the presence of Sox9 suggesting Sox9 activity was mediated within these base pairs. A 54bp S9RE (GRCmm39 19:32,583,940-32,583,993) was also tested throughout this study and was shown to behave similarly to the 32bp S9RE (data not shown).

**Figure 1.**
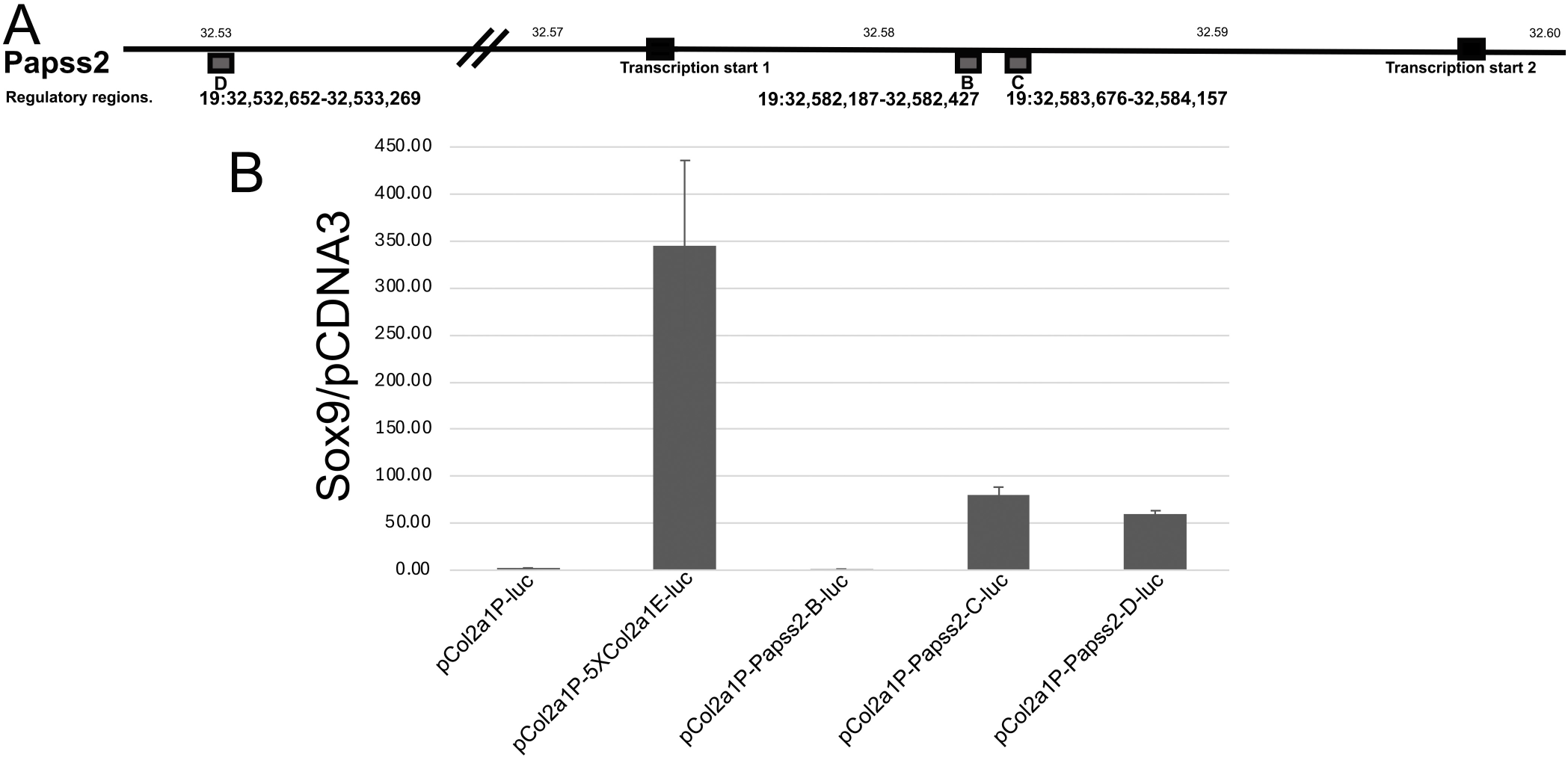
Identification of Sox9 responsive regions near mouse Papss2 gene. (A) <u>Evolutionary conserved regions in and around the mouse Papss2 gene were cloned into luciferase expression vectors.</u> Regions B and C were located in the first intron between the first and second transcription start sites. Region D was located upstream of the Papss2 gene. Gene assembly GRCmm39. (B) <u>Luciferase reporter assays to identify Sox9 responsive regions near Papss2.</u> ATDC5 cells were co-transfected with an expression vector for Sox9 or the control pCDNA3 vector, a renilla luciferase expressing control vector and the indicated firefly luciferase reporter: pCol2a1P-5XCol2a1E-luc is the positive control containing the *Col2a* promoter (Col2a1P-luc) and 5 copies of the *Col2a1* enhancer. A reporter containing the *Col2a* promoter alone (Col2a1P-luc) was used to measure background activity. pCol2a1P-B-luc, pCol2a1P-C-luc, pCol2a1P-D-luc are reporter vectors with each potential enhancer region controlling the Col2a promoter. Renilla luciferase activity was used to normalize luciferase activity which is shown as the ratio of activity from Sox9 expressing cells over that for pCDNA3 expressing cells.

**Figure 2.**
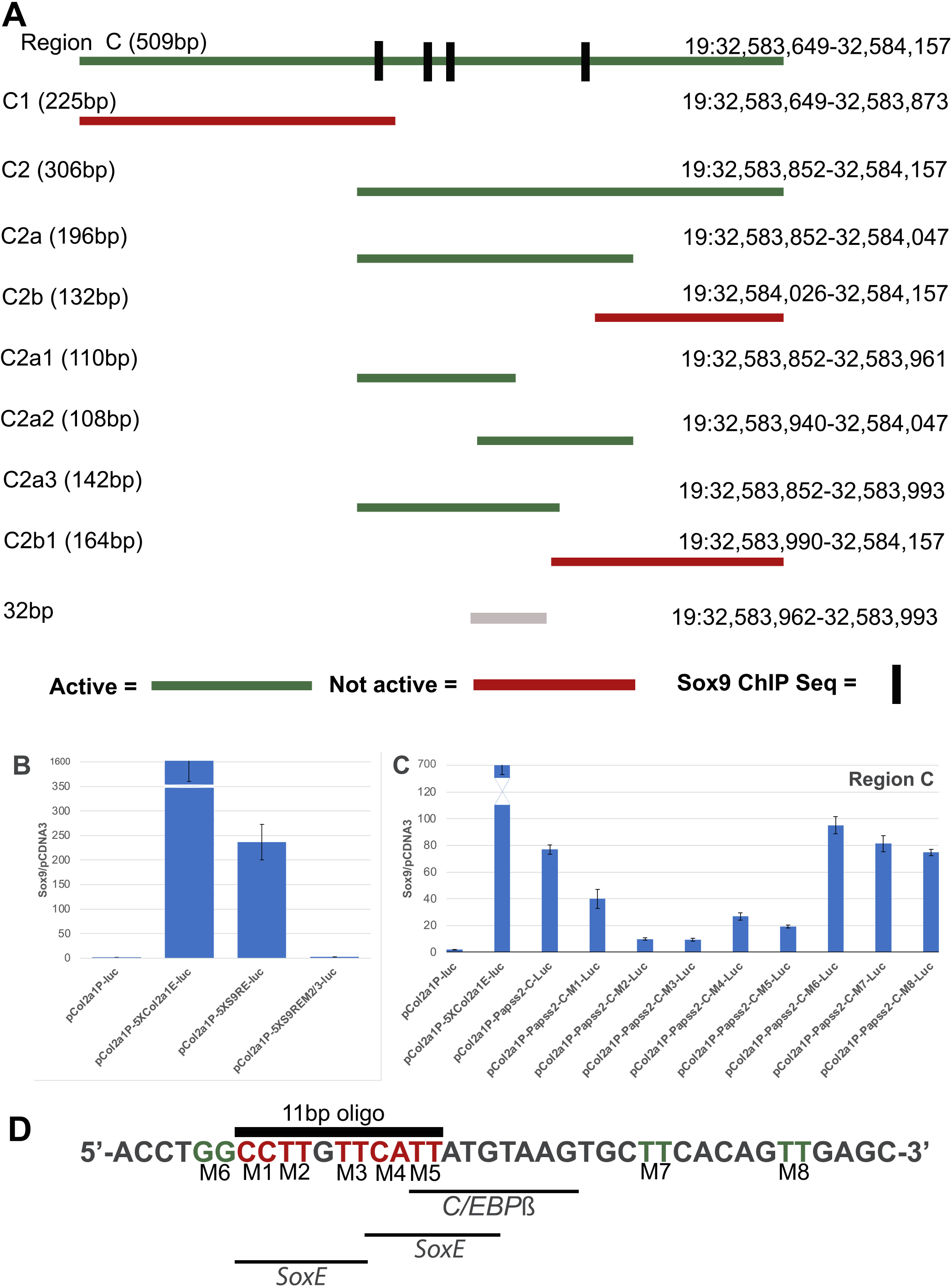
Identification of Papss2 S9RE. (A) <u>Summary of results from luciferase assays focusing on Region C.</u> Green bars represent Sox9 responsive regions. Red represents regions that are not responsive to Sox9 co-expression. Chromosome locations from GRCmm39 assembly are shown. Thick lines in Region C represent locations of Sox9 ChIP Seq peaks from previous studies. (B) <u>Activity of 32bp S9BE.</u> Cells were co-transfected with an expression vector containing Sox9 or a control vector, pCDNA3, and firefly luciferase reporter constructs containing the *Col2a* promoter (Col2a1P-luc) and 5 copies of the *Col2a1* enhancer (pCol2a1P-5XCol2a1E-luc), 5 copies of the *Papss2* S9RE (pCol2a1P-5XS9RE-luc), or mutated S9RE (pCol2a1P-5Xs9REM2/3-luc). The mutation in S9RE contained 4 nucleotide mutation corresponding to M2 and M3 in Figure2D. Co-transfection with a renilla luciferase reporter was used to normalize the data. Results are shown as the average of the firefly/renilla ratio for the Sox9 transfected sample over the firefly/renilla ratio of pCDNA3 transfected samples. Each of three biological experiment was performed in triplicate. A representative experiment is shown. Error bars represent the standard deviation from the experimental triplicate. (C) <u>Luciferase activity for *Papss2* Region C and mutations within the S9RE.</u> Cells were co-transfected with an expression vector containing Sox9 or pCDNA3 and luciferase reporter constructs containing the *Col2a* promoter (Col2a1P-luc) and *Col2a1* enhancer (pCol2a1P-5XCol2a1E-luc), Papss2 Region C (pCol2a1P-C-luc) or Region C containing two nucleotide mutations as shown in Figure 2D (pCol2a1P-C-M1-M5-luc). Co-transfection with a renilla luciferase reporter was used to normalize the data. Results are shown as the average of the firefly/renilla ratio for the Sox9 transfected sample over the ratio for the pCDNA3 transfected samples. Each of three biological experiment was performed in triplicate. A representative experiment is shown. Error bars represent the standard deviation from the experimental triplicate. (D) <u>Sequence of *Papss2*</u> <u>S9RE</u>. Predicted SoxE and C/EBPβ binding sites underlined. Two nucleotide mutations tested that block Sox9 activity are in red (M1-M5). Mutations that did not affect Sox9 activity in green (M6-M8). Location of an 11 bp oligo encompassing most of the predicted SoxE binding sites is shown with a thick black line.

### C/EBPβ suppresses Sox9-mediated Region C activity

C/EBPβ is a known repressor of cartilage enriched genes, including Col2a1 and CD-rap (13–16). To determine if C/EBPβ also acted as a repressor for Papss2, we co-transfected cells with the Papss2 Region C-luciferase reporter, a constant amount of a Sox9 expression vector, and varying concentrations of a full-length C/EBPβ expression vector (Figure 3A). Sox9-mediated luciferase activity was decreased with increasing amounts of C/EBPβ. In contrast, when C/EBPβ levels were held constant and Sox9 levels were increased, there was a dose dependent increase in luciferase activity (Figure 3B). Together the results suggest that the balance of Sox9 and C/EBPβ can regulate Papss2 expression.

**Figure 3.**
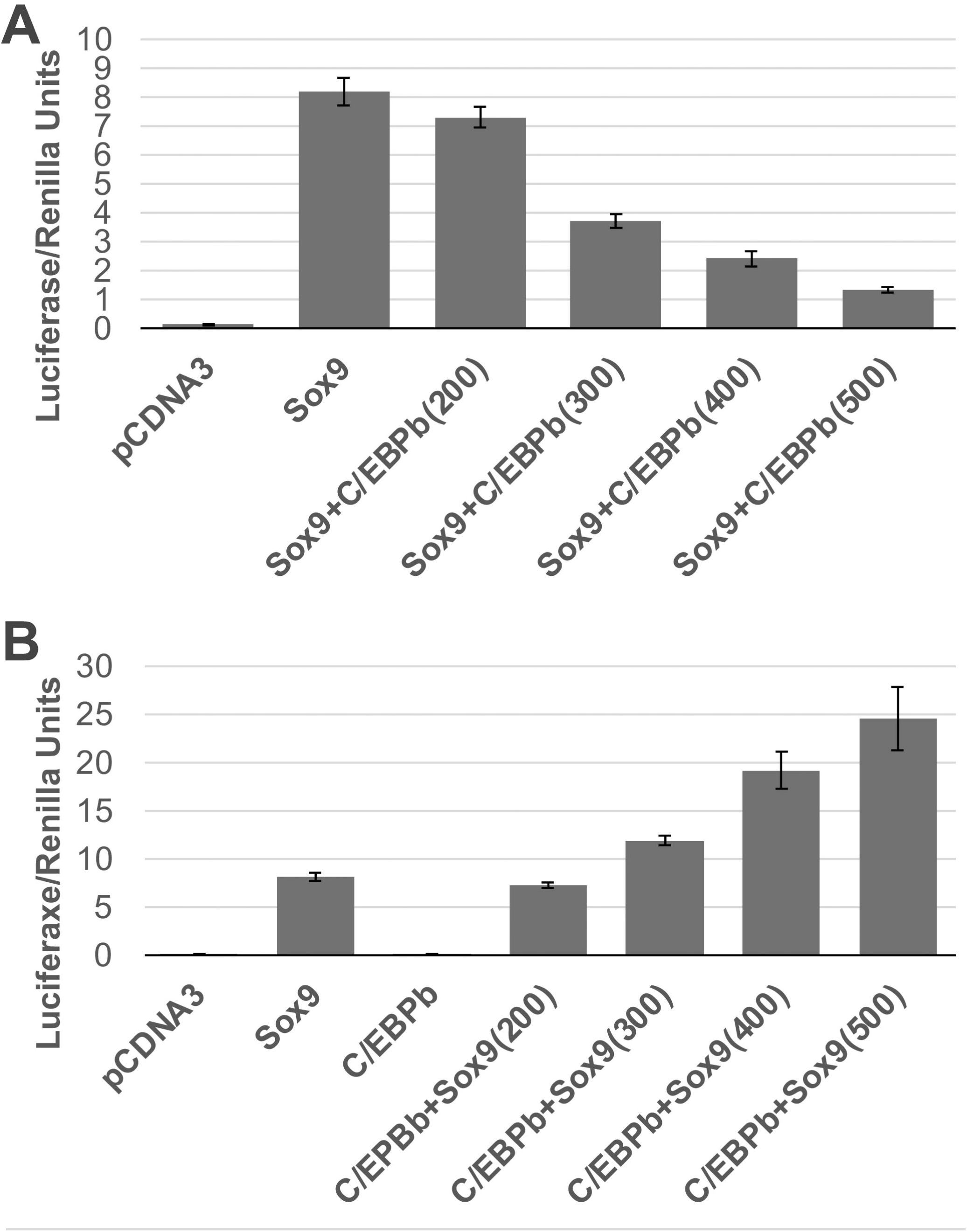
Sox9 and C/EBPβ reciprocally regulate Papss2 S9RE activity. (A) <u>C/EBPβ represses Sox9-mediated transcriptional activity.</u> Cells were co-transfected with the *Papss2* Region C firefly luciferase reporter (pCol2a1P-C-luc) and pCDNA3 or 200ng of the Sox9 expression vector along with indicated amounts of a C/EBPβ expression vector plus pCDNA3 so that the amount of DNA transfected into each cell was equivalent. Co-transfection with a renilla luciferase reporter was used to normalize the data. Cells were harvested 48 hours after transfection and firefly and renilla luciferase activity were measured. The data is shown as the average of the firefly/renilla ratio for each sample. Two separate biological experiments were performed in triplicate. A representative experiment is shown. Error bars represent the standard deviation. (B) <u>Excess Sox9 activates *Papss2* S9RE in the presence of C/EBPβ.</u> Cells were co-transfected with the *Papss2* Region C luciferase reporter (pCol2a1P-C-luc) and pCDNA3 or 200ng of the C/EBPβ expression vector along with indicated amounts of the Sox9 expression vector plus pCDNA3 so that the amount of DNA transfected into each cell was equivalent. Co-transfection with a renilla luciferase reporter was used to normalize the data. Cells were harvested 48 hours after transfection and firefly and renilla luciferase activity were measured. The data is shown as the average of the firefly/renilla ratio for each sample. Two biological experiments were performed in triplicate. Error bars represent the standard deviation.

### Protein binding to the Papss2 S9RE

To test for protein binding to the S9RE, we generated a 40bp DNA probe containing the 32bp S9RE (Figure 1D) to use in Electromobility Shift Assays (EMSA). When the biotin labeled probe was incubated with nuclear extracts from ATDC5 cells under basal conditions, three protein complexes (1-3 from top to bottom) were identified (Figure 4A). The complexes were specific for ATDC5 cells as incubation with nuclear extracts from 293 cells resulted in only one complex that was a different size than those generated with ATDC5 cells (Figure 4A). Complexes were competed away with excess unlabeled 40bp S9RE DNA probe suggesting specificity (Figure 4B). Excess unlabeled probe containing a mutation (M3, Figure 2D) in the predicted SoxE binding site also competed with the formation of protein-DNA complex on the labeled control probe suggesting the mutated site was not required for protein binding to the DNA (Figure 4C). Furthermore, when excess of an unlabeled 11bp oligo overlapping the predicted SoxE binding site (Figure 2D) was incubated with ATDC5 nuclear extracts and the labeled 40bp DNA probe, all three protein complexes remained intact (Figure 4D). There was no competition with the 11bp probe suggesting that the protein complexes observed under basal conditions did not bind to that 11bp of DNA. The results suggest that this region is not necessary for protein-DNA complex formation under basal conditions.

**Figure 4.**
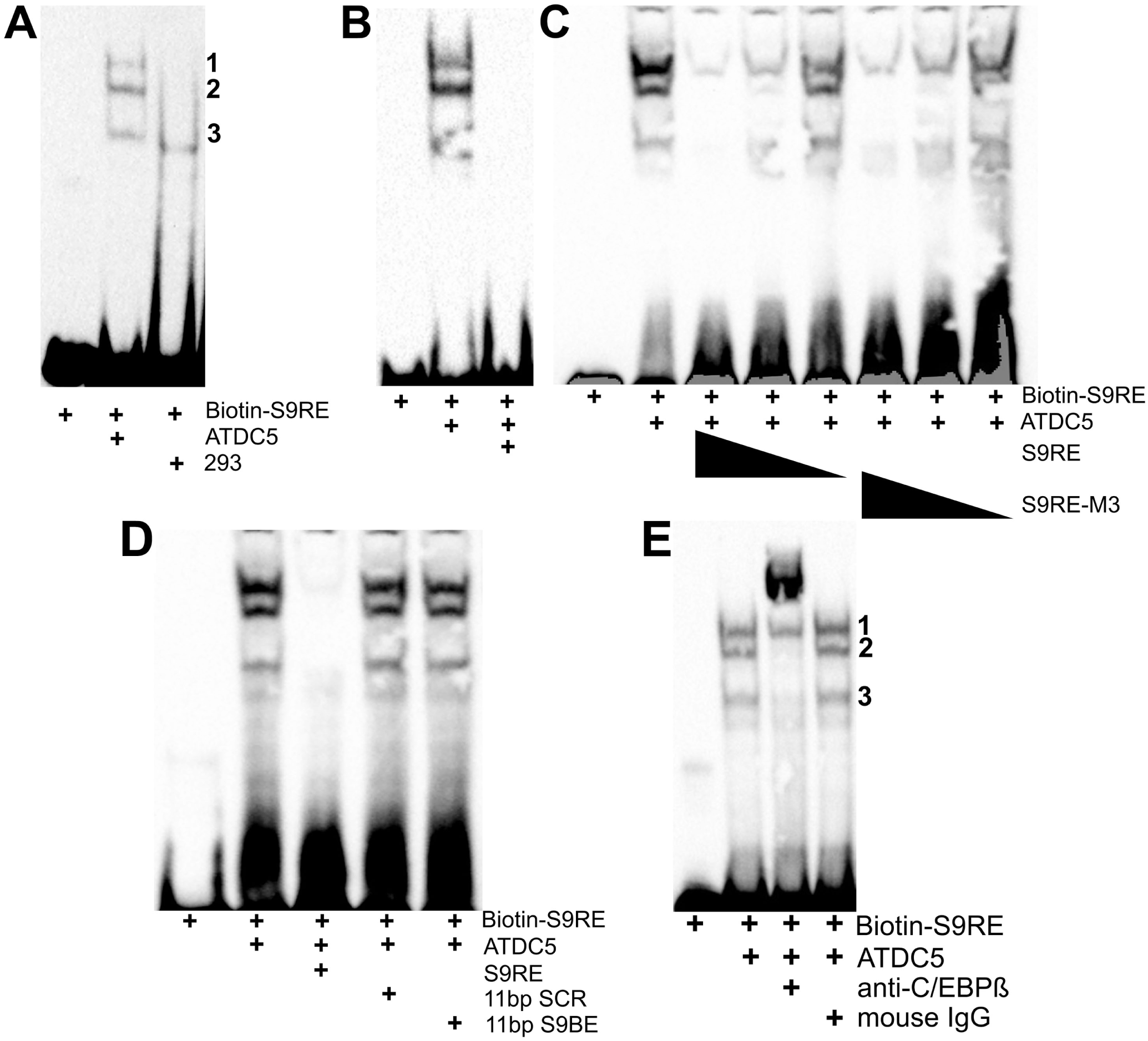
Protein binding to S9RE under basal conditions. (A) <u>Nuclear extracts from ATDC5 cells form three protein complexes on S9RE.</u> A biotinylated 40bp probe shown in Figure 2D (Biotin-S9RE) was incubated with buffer or nuclear extract from ATDC5 cells. Samples were run on a non-denaturing gel, transferred to a nylon membrane and biotin was detected on the membrane using a colorimetric assay. Three bands representing three different protein complexes were observed (1–3). n=14 Only one protein complex with a distinct mobility was observed in samples incubated with 293 cells. n=1 (B) <u>Excess unlabeled S9RE competed with Biotin-S9RE for proteins in the ATDC5 nuclear extract</u> resulting in no protein complexes being observed on the membrane. n=14 (C) <u>Mutations in the predicted Sox9E binding site do not affect protein complex formation on the S9RE.</u> Nuclear extracts from ATDC5 cells were incubated with Biotin-S9RE or Biotin-S9RE and excess unlabeled S9RE at varying concentrations or Biotin-S9RE and excess unlabeled S9RE containing mutation M3 shown in Figure 2D at varying concentrations. n=2 (D) <u>11bp corresponding to a predicted SoxE binding motif does not compete with protein complex formation</u>. Nuclear extracts from ATDC5 cells were incubated with Biotin-S9RE or Biotin-S9RE and excess unlabeled S9RE, a random 11 bp oligonucleotide, or an 11bp oligonucleotide corresponding to the predicted S9BE shown in Figure 2D. n=2 (E) <u>C/EBPβ participates in protein complex formation on the Papss2 S9RE.</u> Nuclear extracts from ATDC5 cells were incubated with Biotin-S9RE or Biotin-S9RE and anti-C/EBPβ antibody or Biotin-S9RE and preimmune mouse IgG. Samples were run on a non-denaturing gel, transferred to a nylon membrane and biotin was detected on the membrane using a colorimetric assay. Samples incubated with anti-C/EBPβ antibody demonstrated a large protein complex at the top of the gel indicating a supershift of the protein complex in the presence of the antibody. n=6

**Figure 5.**
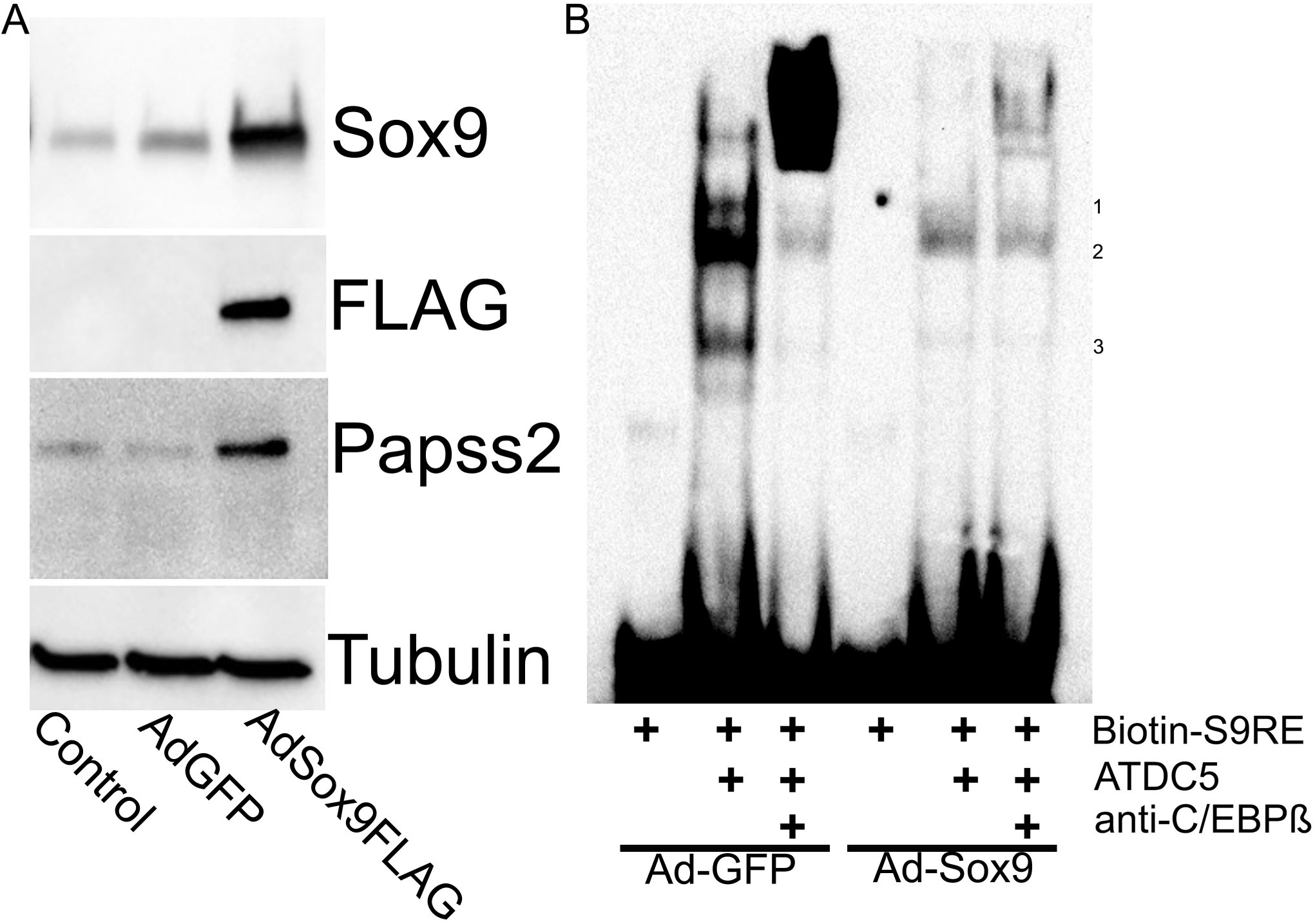
Sox9 regulates protein binding to the S9RE. (A) <u>Western blot confirming Sox9-FLAG expression in ATDC5 cells.</u> Cells were left uninfected (control) or infected with Ad-GFP or Ad-Sox9-FLAG for 48 hours. Protein was isolated and western blot using antibodies to Sox9, FLAG and Papss2 was used to demonstrated relative levels of each protein in the control and infected cells. Alpha-Tubulin was used as the loading control. n=2. <u>(B) Exogenous Sox9 blocks protein binding to the Papss2 S9RE.</u> Cells were infected with Ad-GFP or Ad-Sox9-FLAG. Nuclear extracts were generated and incubated with biotinylated S9RE or nuclear extracts that had been preincubated with antibody to C/EBPβ. n=2

### C/EBPβ binds to Papss2 S9RE

Next, supershift assay was used to determine if C/EBPβ was present in any of the DNA-protein complexes that had been observed in EMSA. When C/EBPβ antibody was incubated with nuclear extract from ATDC5 cells and the labelled 40bp probe, a clear super shift was detected, indicated by the formation of a large protein complex at the top of the gel (Figure 4E). The super shift was specific, control IgG antibodies did not affect formation of any of the DNA-protein complexes observed. The results indicate that protein-DNA complexes on the Papss2 S9RE contain C/EBPβ under basal conditions when Papss2 expression is low.

### Sox9 regulates protein-DNA complex formation on Papss2 S9RE

To determine if Sox9 regulates protein-DNA complexes on the Papss2 S9RE, cells were infected with an Adenovirus containing Sox9 with a FLAG tag (Ad-Sox9-FLAG) or a control Adenovirus (Ad-GFP) as previously described (10). The relative levels of Sox9-FLAG and Papss2 protein were determined by western blot (Figure 6A). Infection with Ad-Sox9-FLAG resulted in high levels of Sox9-FLAG protein expression and up-regulation of Papss2. Equal concentrations of nuclear extracts from Ad-GFP or Ad-Sox9-FLAG cells were used in EMSA assays with the labelled 40bp S9RE probe (Figure 6B). As expected, nuclear extracts from cells that were infected with the control virus generated three protein complexes on labelled S9RE DNA and pretreatment with antibody to C/EBPβ resulted in supershift of the protein complexes. In contrast, nuclear extracts from cells infected with Ad-Sox9-FLAG demonstrated a reduction in overall protein-DNA complex formation relative to Ad-GFP infected controls. The results suggest that Sox9 competes with the formation of protein-DNA complexes on S9RE and may act to derepress Papss2 expression by displacing C/EBPβ.

**Figure 6.**
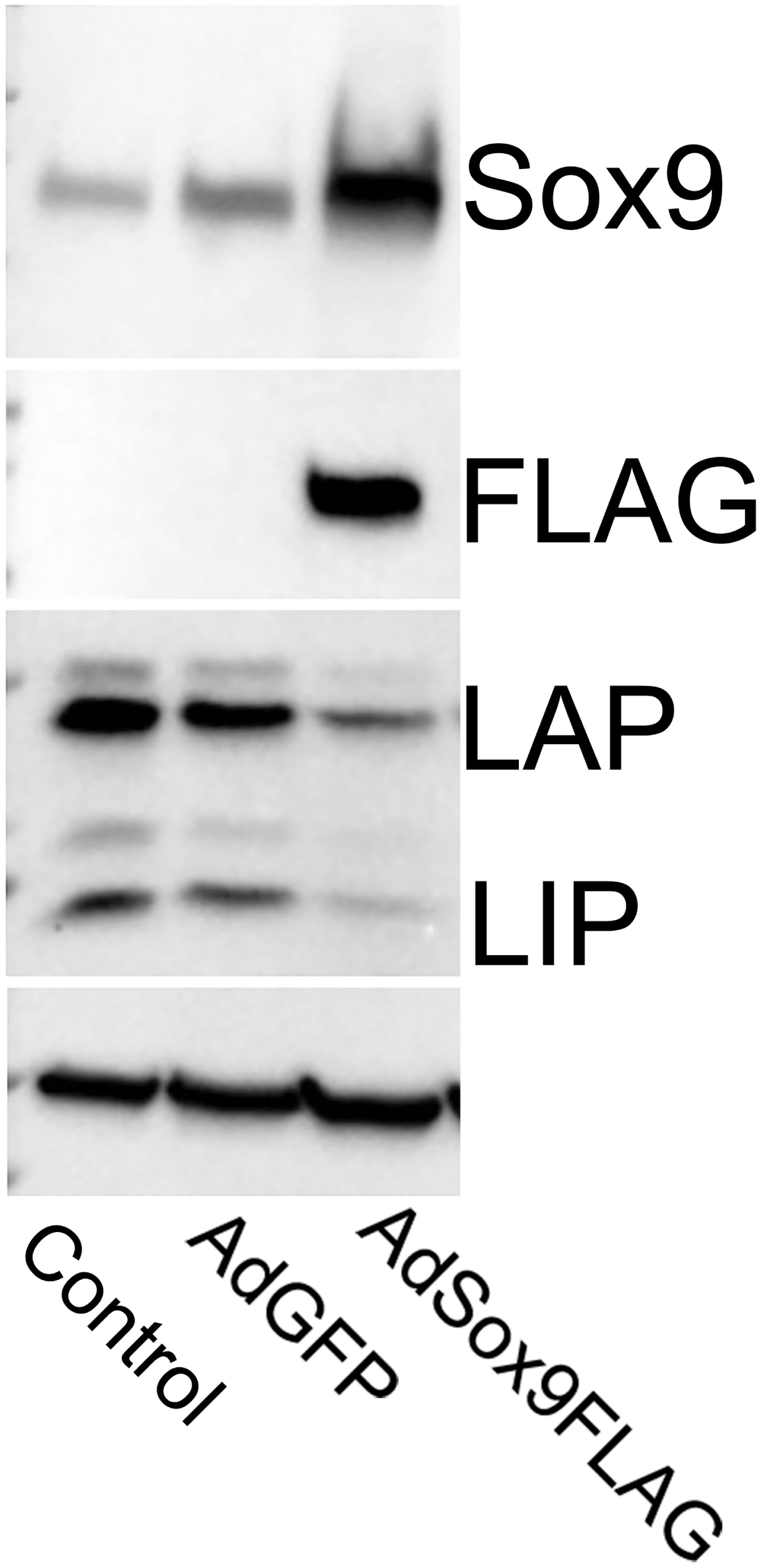
Sox9 down-regulates C/EBPβ protein levels in ATDC5 cells. ATDC5 cells were uninfected (control) or infected with Ad-Sox9-FLAG or Ad-GFP for 48 hours. Protein lysates were generated and Western blot was used to determine the relative levels of C/EBPβ protein, including the LAP and LIP isoforms. Overexpression of Sox9 was also confirmed by western blot using anti-Sox9 and anti-FLAG antibodies. Alpha-Tubulin (a-Tub) was used as a loading control. n=2.

### Sox9 down-regulates C/EBPβ expression and interacts with C/EBPβ protein

Next, the hypothesis that Sox9 regulates expression of C/EBPβ was tested. ATDC5 cells were infected with Ad-Sox9-FLAG or Ad-GFP. Protein expression was determined by western blot (Figure 6). Cells infected with Ad-Sox9-FLAG expressed high levels of Sox9 relative to control and Ad-GFP infected cells. Full-length, LAP, and LIP protein isoforms of C/EBPβ (23) were detected in control and Ad-GFP infected cells. All three isoforms of C/EBPβ protein were reduced in the presence of exogenous Sox9 expression suggesting Sox9 could act in part by down-regulating C/EBPβ protein levels.

The hypothesis that Sox9 protein associates with C-EBPβ protein was tested next using co-immunoprecipitation followed by western blot (Figure 7). ATDC cells were infected with Ad-Sox9-FLAG. Nuclear extracts were generated and Sox9-FLAG was immunoprecipitated using anti-FLAG antibodies. Immunoprecipitation with non-immune IgG was used as a negative control. Immunoprecipitated proteins were separated on a polyacrylamide gel, blotted and incubated with anti-Sox9 or anti-C/EBPβ antibodies. Sox9-FLAG was pulled down with the FLAG antibody and detected on the western blot indicating that the immunoprecipitation worked. Non-immune IgG did not pull down Sox9-FLAG. Primarily, the LAP isoform of C/EBPβ was pulled down in the immunoprecipitation with Sox9-FLAG while the LIP isoform was not detected. We propose that Sox9 acts to derepress Papss2 expression on the 32bp S9RE through down-regulation of expression and direct binding to C/EBPβ.

**Figure 7.**
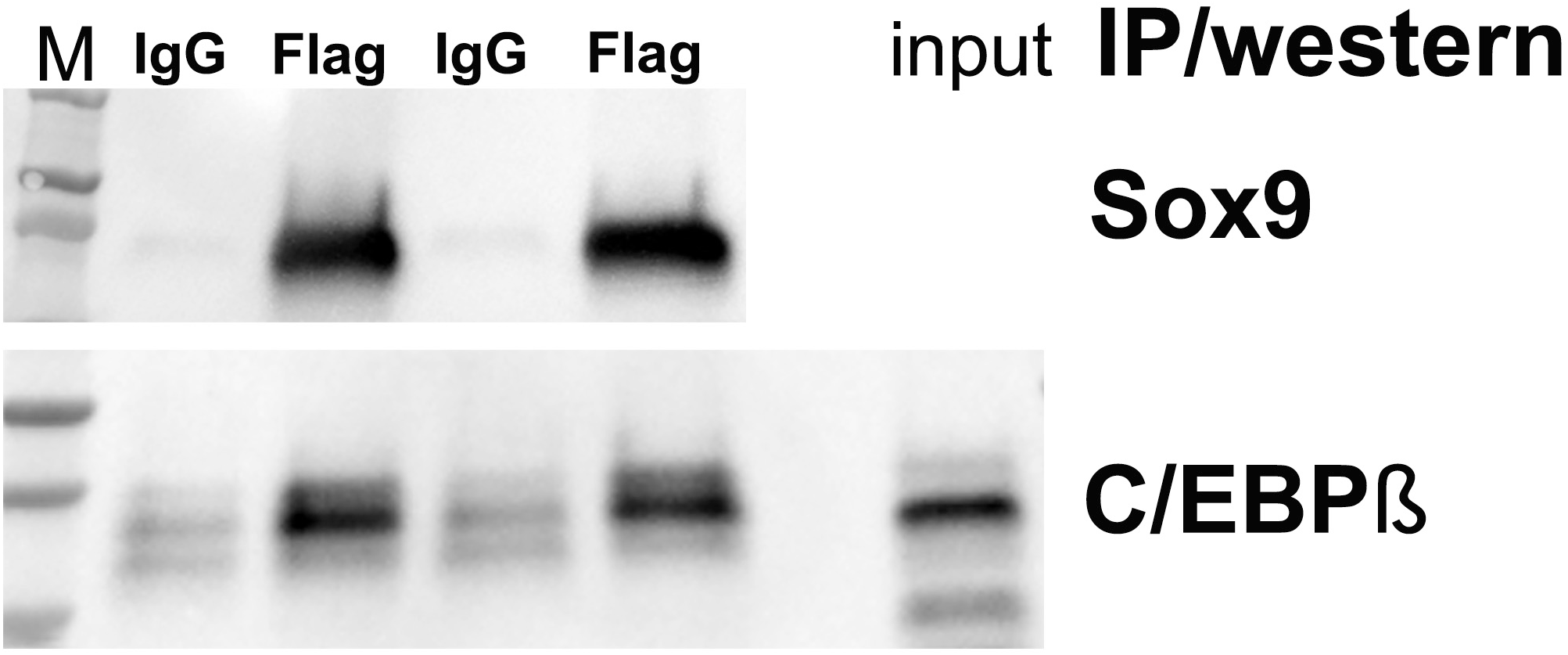
Sox9 interacts with C/EBPβ protein. Cells were infected with Ad-Sox9-FLAG for 48 hours. Cell lysates were immunoprecipitated with anti-FLAG antibody to pull down Sox9 and associated proteins. Immunoprecipitation with a non-immune IgG was used as a negative control. Immunoprecipitated proteins and total input protein was used in western blots to detect Sox9 and C/EBPβ proteins. n=3

## Discussion

Here we identify a 32bp S9RE in the mouse Papss2 gene, located in the first intron upstream of the second start site. Point mutations in a predicted SoxE binding site in the S9RE reduced its activity in response to Sox9. C/EBPβ, a transcriptional repressor, bound to the S9RE under basal conditions. Under conditions of high Sox9 expression, protein binding to the S9RE and total cellular C/EBPβ protein levels were reduced. Sox9 and C/EBPβ were also shown to physically interact. We propose that Sox9 acts to derepress Papss2 transcription through its actions on C/EBPβ on the identified S9RE.

C/EBPβ was previously shown to represses transcription of two cartilage enriched genes, Cd-rap and Col2a1 (13–16). Transcription of Cd-rap requires derepression of C/EBPβ through binding and sequestration by CBP/p300 (site 1; (13, 15). In addition, Sox9 bound to a paired binding site separate from the C/EBPβ site and transcriptional activity of Sox9 was enhanced by CBP/P300 (site 3; (13, 15). Like Cd-rap and Col2a genes, Papss2 S9RE contains a C/EBPβ binding site and co-transfection with C/EBPβ represses Sox9-induced luciferase expression indicating that C/EBPβ is also a repressor for the Papss2 gene. We did not examine the role of CBP/p300 in this study; however, exogenous expression of Sox9 resulted in altered DNA binding to the S9RE in EMSA assays and reduced overall C/EBPβ protein levels. We also found that Sox9 and C/EBPβ proteins interacted using Co-IP. We propose that Sox9 derepresses transcription of Papss2 initially by binding to and sequestering C/EBPβ from the DNA, similar to the roll of CBP/p300 in derepression of Cd-rap, and then down-regulation of C/EBPβ protein acts to maintain Papss2 expression.

Datasets from several ChIP-seq experiments using mature and embryonic cartilage (20, 21) indicate that Sox9 binds somewhere within a 170bp to 550bp region that includes the 32bp Papss2 S9RE. However, the peak of the sequence alignments were not within the responsive element we identified, the closest was 30 bp upstream in Region C2a1 (Figure 2). A 227bp CBP/p300 binding region that surrounds the Papss2 Sox9 response element was also identified in cartilage by ChIP-seq (20). The ChIP-seq peak alignment for CBP/p300 is within a few base pairs of the Sox9 binding peaks in Region C2a1. It is possible that this region is analogous to the Sox9 binding region in Cd-rap (site 3) and the 32 bp C/EBPβ binding region we identified here is similar to site 1 (14, 15). ChIP seq data for C/EBPβ is not available at this time so we don’t know if there are other C/EBPβ binding sites within Region C.

Attempts to detect Sox9 binding to the 40- and 54bp Papss2 S9RE using ChIP or EMSA supershift assays were inconclusive. There could be several reasons for this including technical issues with the antibodies used. In addition, the SoxE binding sites identified in the 32bp S9RE have a weak prediction score and more closely aligned with Sox8 binding motifs in the Jaspar 2024 database (22). Sox9 and Sox8 are both members of the SoxE subgroup within the Sox family of HMG-box type DNA binding proteins (7). SoxE proteins, Sox8, Sox9, and Sox10 are expressed in an overlapping manner in several tissues. Sox8 and Sox9 have some overlapping activities in the skeleton (8). Sox8 and Sox9 both promote skeletal development by increasing chondrocyte proliferation and differentiaion. Overexpression of either Sox8 or Sox9 in chondrocytes promotes cell proliferation, even when the other Sox protein is inactivated (8). It is possible another SoxE protein binds to this site or that binding is so weak that it cannot be detected by conventional methods.

Alternatively, other proteins may bind to this region and directly or indirectly mediate Sox9 activity. There is a peak ChIP seq binding site recorded for Nr3c1 within the 32bp S9RE overlapping with sequences that are involved in Sox9-activity (24). Nr3c1 (nuclear receptor subfamily group C member 1 protein) is a glucocorticoid receptor that acts a transcription factor when activated (25). Variants in human NR3C1 are associated with osteoarthritis (26). There are also sequences with high Jaspar 2024 prediction scores for other transcription factors in this region including NR5A2 (score 402) and PouF1 (Score 404). Nr5a2 is a DNA-binding zinc finger transcription factor and a member of the fushi tarazu orphan nuclear receptor family. It has been shown to promote diverse connective tissue fates in zebrafish jaw (27). PouF1/Brn3 is a member of the POU-IV class of neural transcription factors and not know to be expressed in cartilage (28).

In summary, we have identified a Sox9 responsive DNA element in the first intron of mouse Papss 2 that appears to regulate Papss2 through Sox-9 medicated derepression of C/EBPβ.

## Materials and Methods

### Cell culture

ATDC5 cells (29) were suspended and cultured in Dulbecco’s Modified Eagle Medium/F12 medium (DMEM/F12, Life Technologies Corp, # 11320-033) containing 5% heat-inactivated fetal bovine serum (FBS, Fisher Scientific), 1% penicillin-streptomycin (Life Technologies Corp, # 15140-122), and 1% L-glutamine (Life Technologies Corp, # 25030-081). Cells were grown in 5% CO2 at 37°C in humidified incubator. The cells were passaged at 1:6 after confluence.

293A cells were cultured with Dulbecco’s Modified Eagle Medium (DMEM, Life Technologies Corp, # 11995-065) supplemented with 10% heat-inactivated fetal bovine serum, 1% penicillin-streptomycin, and 1% L-glutamine.

### Construction of luciferase reporter plasmids

A Col2a1 luciferase reporter plasmid used as a positive control for Sox9 activity, pCol2a1-5x-Luc, was obtained from Dr. Veronique Lefebvre (18). The vector contains 5 repeat Sox9-reponsive enhancers specific for the *Col2a1* gene upstream of the minimum 95bp Col2a1 promoter. A new luciferase reporter, in which the enhancers were removed and the 95bp Col2a1 promoter retained, was generated to test potential enhancers from the *Papss2* gene. This vector was referred to as pCol2a1P-Luc.

Primers specific for regions of the mouse *Papss2* gene were designed using Snapgene software. The clone from the Bac DNA RP23 library containing the *Papss2* gene was obtained from BACPAC Resource Center (Oakland, CA, USA) and was used as the template to amplify DNA fragments from the *Papss2* gene. These PCR fragments were digested and inserted upstream of 95bp minimal promoter in pCol2a1promoterLuc.

The QuikChange Lightning Site-Directed Mutagenesis Kit, (Cat. Number: 210518, from Agilent Technologies) was used for all DNA mutagenesis.

### Cell Transfection

ATDC5 cells were plated in 24-well plates at 2X10^5^ cells/well in 1ml of culture medium and cultured for overnight. The cells were grown to 80-90% confluence before transfection. The cell transfection was performed as described in the instructions for the ViaFect reagent kit (Promega, Corp). Briefly, 300ng of luciferase reporter plasmid, 30ng of plasmid containing Renilla Luciferase (to control for transfection efficiency; Promega, Corp1), and 200ng of relevant expression or control plasmid were suspended and mixed in Opti-MEM (Life Technologies Corp). Next, 3.5ul of Viafect transfection reagent was added to a total volume of 50ul. The plasmids and Viafect were mixed and incubated at room temperature. After 20-30 minutes, the mixture was dropped to wells and the cells were cultured for 24 hours before luciferase assays were performed.

### Luciferase assay

Luciferase activity was measured according to the method included in the Dual-Luciferase Reporter Assay kit (Promega, Corp). Briefly, the culture medium was removed from the plate and cells were washed once with PBS. Then 300ul of 1X Passive Lysis Buffer was added on top of the cells and the plate was incubated at room temperature for 15 minutes while shaking. The cell lysis sample was mixed by pipetting and transferred to an Eppendorf tube. Debris was spun down and the supernatant was recovered for luciferase assay or stored at - 20C.

For the luciferase assay itself, 10ul of cell lysis sample was mixed with 50ul of substrate. The luciferase activities were then determined with 20/20ª Luminometer (Turner Biosystems). After that 50ul of stop solution was added into the above testing sample. After mixing the activity for Renilla luciferase was determined using the Luminometer.

### Electrophoretic Mobility Shift Assays

A Nuclear Extract Kit (Active Motif North America, # 40010) was used for preparation of nuclear proteins. Briefly, cells grown to confluence in a 100 mm tissue culture plate were washed with ice-cold PBS/phosphatase inhibitors. The cells were gently suspended in 500ul of 1X Hypotonic Buffer and incubated for 15 minutes on ice for swelling. After addition of 25ul detergent the cells were vortexed for 10 seconds at the highest setting. After spin-down the ATDC5 nuclear pellets were recovered. Next, 200ul of complete lysis buffer was added to the recovered nuclei and the pellets were homogenized for 5 minutes. The supernatant (nuclear fraction) was stored at -80C.

EMSA were performed as described in the Gelshift Chemiluminescent EMSA kit (Active Motif North America, # 37341). Double-stranded biotin 5’ end-labeled and unlabeled oligonucleotides used as probes were synthesized by IDT (Integrated DNA Technologies). Briefly, 5ul of nuclear extract was incubated with the relevant biotin 5’ end-labeled DNA probe. Samples were then resolved by electrophoresis on a 6% DNA Retardation Gel (Life Technologies, # EC6365BOX) and transferred to a positively charged nylon membrane. The biotin end-labeled DNA probe was detected using streptavidin conjugated to horseradish peroxidase (HRP) and a chemiluminescent substrate. For competition experiments, different amounts of unlabeled DNA probe were incubated with nuclear extracts. For super shift assays, C/EBPβ antibody (Cat # Santa Cruz Biotechnology, # sc-7962, raised in mouse, monoclonal) and control isotype antibody (Santa Cruz Biotechnology, # sc-2025) were added and incubated with nuclear extracts for 1 hour at 4C before EMSA was performed.

### Adenovirus infection

Adenovirus encoding wild-type, FLAG-tagged SOX9 with IRES regulated enhanced green fluorescent protein (eGFP) (Ad-SOX9) and control virus encoding for only eGFP (Ad-eGFP) were used in this study (10). ATDC5 cells were cultured overnight and infected with adenoviruses by adding 75 M.O.I. of adenovirus into culture media. After 48h of culture, the cells were washed with PBS and lysed with immunoprecipitation (IP) buffer (Cell Signaling Technology, # 9803) for Co-IP western blot assay or the cells were used for nuclear extract preparation for EMSAs.

### Western blot

Fifty μg of protein lysate per sample was separated by reducing electrophoresis on 4–20% polyacrylamide gels (Bio-Rad Laboratories, # 456–8096). Protein was transferred from gels to polyvinylidene fluoride membranes (Bio-Rad Laboratories, # 162–0177) using a Trans-Blot Turbo Transfer System (Bio-Rad Laboratories, # 1704150). Membranes were blocked with 3-5% milk (Santa Cruz Biotechnology, # sc-2325). Membranes were then incubated overnight with anti-sox9 primary antibody (1:1000, Sigma Aldrich, # AMAB90795, raised in mouse, monoclonal), anti-C/EBPβ primary antibody (1:200, Santa Cruz Biotechnology, # sc-7962, raised in mouse, monoclonal). To assess whether equivalent amounts of protein were loaded in all wells, an anti-cyclophilin B primary antibody (1:1000, Abcam, # ab16045, raised in rabbit, polyclonal) was used. Membranes were washed with Tris-buffered saline containing 0.05% Tween 20 (TBST) and incubated with HRP-conjugated anti-mouse (1:2000, cell signaling Technology, # sc-2055, raised in goat) or anti-rabbit (1:2000, Cell Signaling Technology, # 7074S, raised in goat) secondary antibodies. Images of Western blots were acquired on a ChemiDoc MP system (Bio-Rad Laboratories).

### Co-immunoprecipitation-western blot

For Co-IP-western blot, 300 μg of protein lysate was used for each immunoprecipitation. To reduce nonspecific background, the protein lysates were treated for 3h in the presence of magnetic beads ((Thermo Scientific, #88847) and control normal mouse IgG (Santa Cruz Biotechnology, #sc-2025). The cleared protein lysates were recovered by magnetic separation. Immunoprecipitation was performed by incubating cleared samples with 20 ul of protein G magnetic beads and anti-FLAG primary antibody (5ug, Sigma Aldrich, #F1804, raised in mouse, monoclonal) or normal mouse IgG, as control, overnight. The magnetic beads were isolated and washed three times. Finally, the protein complex was recovered with magnetic beads and 1X protein loading buffer (Cell Signaling Technology, #56036S). Samples were run on a polyacrylamide gel and blotted to polyvinylidene fluoride membra nes. For western blot anti-FLAG primary antibody, anti-Sox9 primary antibody (1:1000, Sigma Aldrich, #AB5535, raised in rabbit) or anti-CEBP beta primary antibody (1:1200, Boster Biological Technology, #M01100, raised in rabbit, monoclonal) were used. The HRP-conjugated anti-rabbit (1:2000, Cell Signaling Technology, # 7074S, raised in goat) secondary antibodies were used as above.

